# The effect of circulation rate on the water quality of container-type recirculation aquaculture system under different feeding rates of *Schizothorax wangchiachii*

**DOI:** 10.1101/2023.06.22.546147

**Authors:** Junming Zhou, Qinglin Cai, Kelei Zhou

## Abstract

As an experimental fish, juvenile *Schizothorax wangchiachii* were fed different dosages (100, 300, or 500 g), and the changes in water quality in the inlet and outlet of the biofilter and in the aquaculture buckets were measured under different circulation rates (1.5, 3, or 4.5 m^3^/h; water samples were collected every 2 hours). With 100 g of feed and different circulation rates (1.5, 3, or 4.5 m^3^/h), the average removal rates of ammonia nitrogen and nitroso nitrogen by the biofilter were 40.4, 36.8, and 27.7% and 43.9, 36.9, and 29.8%, respectively. With 300 g of feed, the average removal rates of ammonia nitrogen and nitroso nitrogen were 42.3, 37.9, and 33.6% and 45.8, 32.7, and 21.7%, respectively. With 500 g of feed, the average removal rates of ammonia nitrogen and nitroso nitrogen by biofilters with different circulation rates were 53.5, 42.6, and 35.7% and 40.3, 26.5, and 18.1%, respectively. Therefore, when the feeding amounts were 100, 300, and 500 g, respectively, the optimal circulation rates were 1.5, 1.5, or 4.5 m^3^/h, which allowed the best management and reduction in energy consumption of the circulating aquaculture system. This research conclusion provides a reliable scientific reference for the promotion of a low-cost new model of industrial circular water aquaculture.

## 1. Introduction

Aquaculture water is mostly sourced from natural water bodies, and thus, a good ecological environment is a fundamental foundation for the sustainable and healthy development of the aquaculture industry (Lithgow *et al*., 2019). Although shrimp farming is one of the pillars of China’s aquaculture industry, the aquaculture technology in some regions has not yet been improved (Patrona, 2006; Xie & Yu, 2007; A Sánchez-Paz & Muhlia, 2020). If traditional aquaculture methods with high density and water exchange rates persist, where some even abuse drugs, the aquaculture wastewater remains untreated. In addition, it is directly discharged into natural waters, causing eutrophication of receiving rivers, lakes, and seawater areas, which is already worsening day by day. The issue of environmental pollution caused by aquaculture has attracted global attention (Naylor *et al*., 2000; Pez-Osuna, 2001). In order to change traditional aquaculture techniques and models, intensive research has been carried out both domestically and internationally on circulating water aquaculture models (Davis & Amold, 1998; Huang *et al*., 2010).

Circulating water fish farming systems apply modern engineering principles and methods to aquaculture engineering (Mingfeng *et al*., 2000; Brown *et al*., 2016), with major characteristics of water-saving, land-saving, high-density intensification, and controllable emissions; in short, they meet the requirements of sustainable development. In recent years, aquaculture has rapidly developed in China, especially with the increase in aquaculture varieties and innovation in water treatment technology (Hixson, 2014). Circulating water fish farming system mainly relies on artificial feeding to promote fish growth and increase aquaculture yield (Zhang *et al*., 2011). Only a portion of the nitrogen input with feed is absorbed by the fish, while the rest enters the water body that receives the aquaculture discharge. Due to the inherent assimilation efficiency of nutrients by fish, coupled with a significant portion of their intake entering the water body in the form of excreta, the nitrogen load in aquaculture increases, and the ecological environment on which fish rely for survival is vulnerable to damage (King & Warburton, 2007).

In recirculating aquaculture systems, the total ammonia nitrogen emissions can be calculated through empirical constants (Khangembam *et al*., 2017). The volume of the biofilter is calculated based on tolerance concentrations of ammonia and nitrite nitrogen and the nitrification rate of the biofilter (Eding *et al*., 2006; Yang *et al*., 2019). However, the ammonia emission rate for farmed animals varies with fish activity; especially after feeding, there is a peak in ammonia nitrogen emissions. If the biological filter is designed based on the average load, it cannot completely remove the ammonia nitrogen during the peak period in a timely manner. This creates the risk of exceeding the acceptable concentration of ammonia nitrogen in the water body. Improving the circulation rate in the water system can significantly reduce the concentration of ammonia and nitrite nitrogen in the, optimize the aquaculture environment, and promote fish growth (Alongi *et al*., 2009). Theoretically increasing the circulation flow during the peak period of ammonia nitrogen can also reduce the mass concentration of ammonia nitrogen. Here, combined with normal flow rate operation at other times, the principle of variable speed flow control technology has been proposed to achieve water quality similar to the constant operation of high circulation flow, in order to reduce operating costs.

*S. wangchiachii* belongs to Cypriniformes, Cyprinidae, Schizothoracinae, and Schizothorax families. It lives in plateau areas and is a small to medium-sized cold-water fish. This species occurs with high abundance in some places, such as the Jinsha River, where it is an economically important species. *S. wangchiachii* has tender flesh, a delicious taste, high protein content, low fat content, and low cholesterol. Due to the demand for low-fat and high-protein foods, *S. wangchiachii* has become a widely popular high-value food (Dai *et al*., 2018; Hou *et al*., 2019; Wu *et al*., 2020).

Based on the study of ammonia excretion patterns and nitrification performance of the *S. wangchiachii* biofilter, this experiment provides a theoretical reference for the design of efficient circulating water biofilters.

## 2. Methods

### 2.1 Experimental design

The feeding conditions for the experimental aquaculture design were two meals per day according to the actual needs of the aquaculture industry. The interval between feeding was 12 hours. It is necessary to adjust the circulation volume to increase the processing capacity of the biofilter and allow the three-state nitrogen indicators in the breeding bucket to reach a safe range. During the 12-hour period, it is necessary to adjust the circulation rate at intervals to control the aquaculture water quality based on the amount of feed and changes in water quality. According to the feeding settings during production, optimally setting the amount of water for circulation in the aquaculture can effectively save breeding costs and help achieve the goal of precise aquaculture. Therefore, under different feeding (100, 300, or 500 g) and circulation rates (1.5, 3, or 4.5 m^3^/h) in the aquaculture mode, the first water sample collection in the experiment was immediately after feeding, and the start time was recorded. Water samples were collected every two hours, and the experiment ended within 12 hours. The experiment was also started under different experimental conditions, whereby water replacement was done, and the circulating water system was started for three hours until the water quality stabilized. It was ensured that there were no other influencing factors that could change the water quality indicators. The starting conditions of the experiment were kept the same, and the concentrations of the indicators at the beginning of the experiment were controlled within a similar range.

### 2.2 Experimental animals and feed

The experimental fish were selected to be *S. wangchiachii*; 200 healthy and similarly weighted *S. wangchiachii* were selected. They were placed in breeding barrels 1 and 2 in equal numbers. After a week of temporary cultivation, the experiment started based on the actual situation. The formal experiment of the circulating water aquaculture system’s biofilter presented a good water treatment effect, indicating that the biofilm pre-cultivation was successful. Table 1 presents values for feed composition analyses. The experimental feed was based on freshwater fish expansion formula feed that consisted of fish meal, soybean meal, peanut meal, flour, soybean oil, corn protein meal, calcium dihydrogen phosphate, zinc sulfate, vitamin A, vitamin C, vitamin D3, and vitamin E folate, etc.

**Table 1.** Guaranteed values of feed composition.

| Number | Component | Values<br>(%) |
| --- | --- | --- |
| 1 | Crude protein | $\geq 43.0$ |
| 2 | Crude fat | $\geq 5.0$ |
| 3 | Coarse ash content | $\leq 15.0$ |
| 4 | Lysine | $\geq 2.20$ |
| 5 | Coarse fiber | $\leq 6.0$ |
| 6 | Water content | $\leq 12.0$ |
| 7 | Total phosphorus | $\geq 1.00$ |
| 8 | Calcium | 1.00~4.00 |
| 9 | Sodium chloride | 0.50~3.00 |

### 2.3 Experimental design

Figure 1 shows the process scheme of the precision production experimental system for industrial circulating water. The circulating water aquaculture system mainly includes water pumps, coarse filter barrels, aquaculture barrels, ultraviolet rays, biological filters, protein separators, etc. The breeding bucket and coarse filter bucket are connected through PVC pipes, and the overall water level remains consistent. When the circulating water pump is turned on and working, the water in the coarse filter barrel can be circulated through the connecting water pipe via control through a computer. The water level in the coarse filter barrel drops, causing a water level difference between the breeding barrel and the coarse filter barrel. A certain water level difference induces pressure, which pushes the water from the breeding bucket into the coarse filter bucket. The circulating water pump forms a water flow through the connecting pipeline; water passes through the cooling and heating machine and then enters the biological filter to return to the breeding bucket through the diversion switch. Another diverged flow of water enters the ultraviolet sterilizer and flows back to the protein separator before returning to the breeding bucket. The return water flow of breeding buckets 1 and 2 was designed based on the limited volume of standard containers and was controlled using PVC ball valves. The experiment was designed to adjust the consistency of the return water flow in breeding buckets 1 and 2, which was equivalent to the consistency of the changes in the aquatic environment of breeding buckets 1 and 2. The goal was to achieve the efficient utilization of limited space and save resources, which is a major feature of containerized recirculating aquaculture systems.

**Fig. 1.**
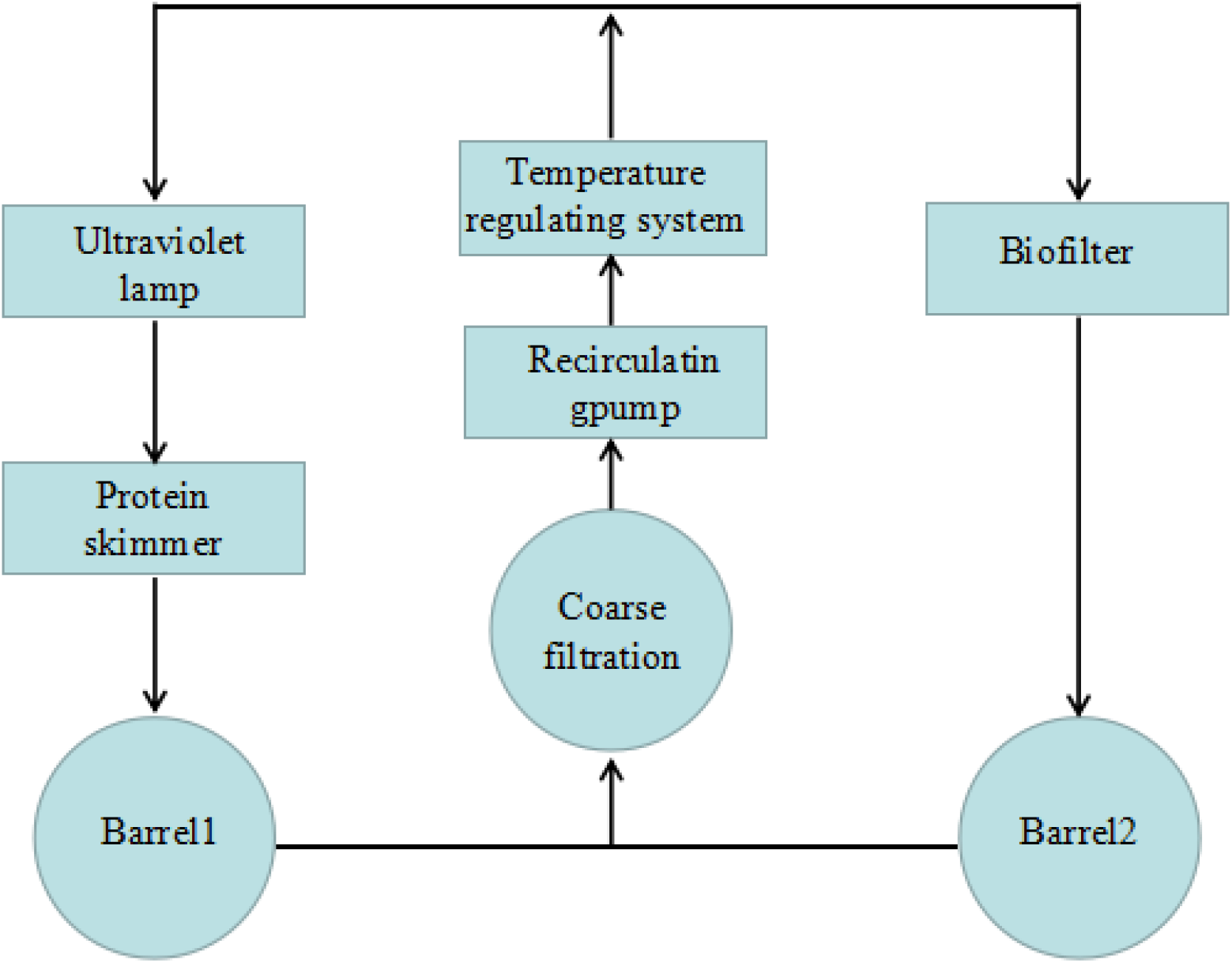
Flow process of factory circulating water in the precision production experimental system.

### 2.4 Breeding management

The experimental fish were selected to be *S. wangchiachii* species, which had been raised by Liangshan Xide Aquatic Co., Ltd., and were temporarily raised for a week before selection. Two-hundred *S. wangchiachii* each were placed in aquaculture buckets 1 and 2, and the circulating water was diverted back to the two aquaculture buckets to regulate uniform and equal water flow. The chiller and heater were maintained at 13°C, which was a good temperature for the growth of *S. wangchiachii*. Environmental parameters that were monitored and recorded daily included temperature, dissolved oxygen, pH, and salinity. Two regular meals were fed every day (at 08:00 and 20:00); the feeding amount was set according to specific experimental conditions.

### 2.5 Water sample collection

The day before the official start of the experiment, the same amount of feed and circulation rate was set, and the experimental fish were adapted to the aquaculture environment. Before the experiment started, the aquaculture water and cycle were changed for about two hours. The first water samples were collected from aquaculture buckets 1 and 2, as well as the inlet and outlet of the biological filter. Immediately after the collection of the first water samples, the feeding amount was set, and the timing for the experiment was started. Water samples were collected every two hours until the last 12-hour collection had been completed. When setting experimental conditions for feed and circulation rates each time, in order to ensure the stability of the initial water quality index concentration, it was deemed necessary to thoroughly replace the water and cycle it for 2 hours at the same circulation rate uniformly. Water samples were then collected after the biofilm was adapted to the new aquatic environment.

### 2.5 Determination of serum and tissue indicators

The method for measuring water quality indicators was as follows: TAN: naphthalene colorimetric method; NO_2_-N: N-1-nitrohexidine photometric method; and NO_3_-N: zinc cadmium reduction diazo azo colorimetric method.

The equation to calculate the percentage removal of pollutant concentrations (referred to as the removal rate) is:

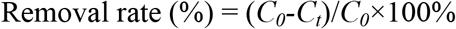

In the formula, *C*_*0*_ and *C*_*t*_ are the pollutant concentrations (mg/L) in the inlet and outlet of the filter, respectively.

The equation to calculate the daily average removal rate by the filter material per unit area (referred to as the removal rate):

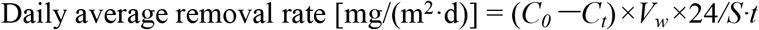

In the formula, *C*_*0*_ is the concentration of pollutants at the inlet of the filter (mg/L); *C*_*t*_ is the concentration of pollutants at the outlet of the filter after retention for of the shrimp pond water time *t* (h); *V*_*w*_ is the water storage capacity of the filter (m^3^), and *S* is the total surface area of the filter material (m^2^).

## 3. Results

### 3.1 The effect of different circulation rates on the water quality of the aquaculture bucket at a feed rate of 100 g

Figure 2a shows the variation in ammonia nitrogen concentration in the aquaculture barrels over time (0–12 hours) at different circulation rates (1.5, 3, or 4.5 m^3^/h) under the initial environmental conditions of 100 g of feed and fresh water. When the circulation rate was adjusted to 1.5, 3.0, or 4.5 m^3^/h, the water treatment system started to work effectively. The ammonia nitrogen concentration in breeding barrels 1 and 2 increased synchronously over time and then decreased. The initial values of ammonia nitrogen concentration in the breeding barrels 1 and 2 at the 0th hour were 0.38 and 0.38 mg/L; 0.36 and 0.37 mg/L; and 0.35 and 0.33 mg/L, respectively: These values indicate that the performance of the biofilter was relatively stable, and the initial conditions set in the experiment were within the error range.

**Fig. 2.**
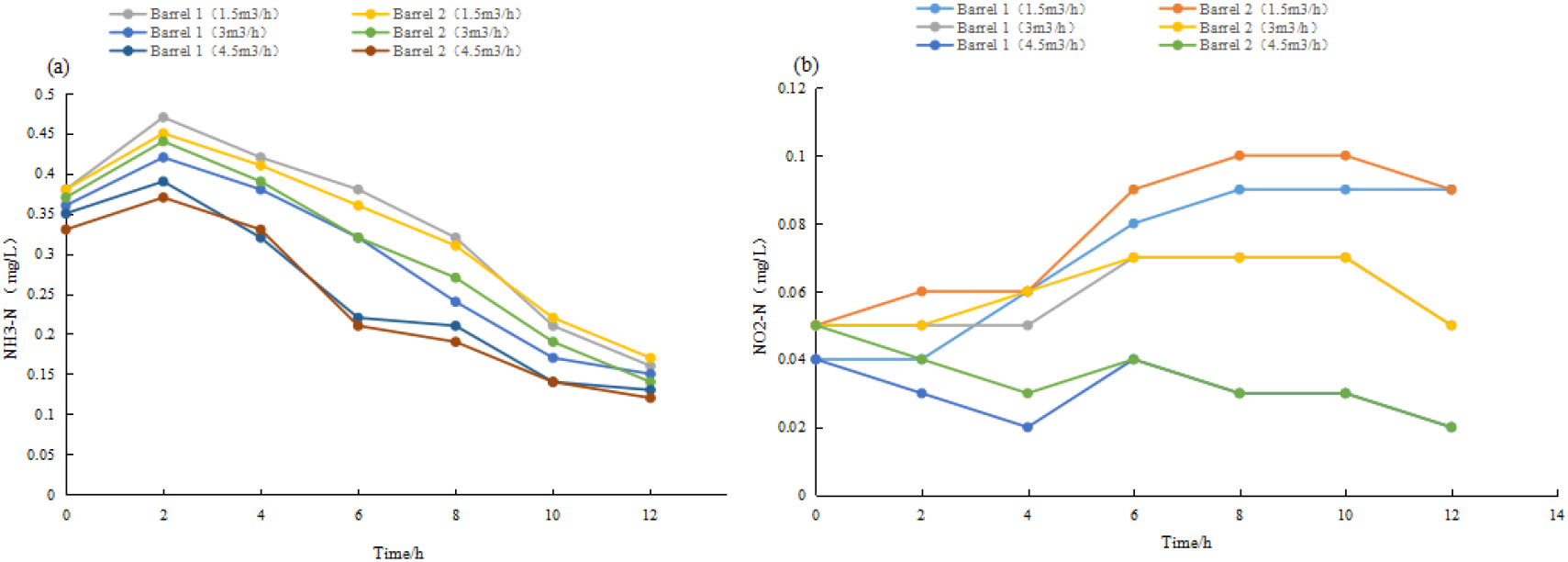
Changes in ammonia nitrogen and nitroso nitrogen in breeding barrels with different cycle rates over time with 100 g of feed.

There was no significant difference between the treatments. During the second hour, the maximum values of 0.47 and 0.45 mg/L; 0.42 and 0.44 mg/L; and 0.39 and 0.37 mg/L were recorded in breeding buckets 1 and 2, respectively. This indicated that the feces produced by the offspring of *S. wangchiachii* entered the water body to accumulate ammonia nitrogen concentration, leading to an increase in ammonia nitrogen concentration. After two hours, the decrease in ammonia nitrogen concentration reflected the gradual, but effective, removal of harmful substances by the biofilter. At the 12th hour, the aquaculture buckets 1 and 2 reached minimum values, i.e., 0.16 and 0.17 mg/L; 0.15 and 0.14 mg/L; and 0.13 and 0.12 mg/L, respectively. At this time, the effectiveness of the biofilter reached the optimal removal level, helping to maintain extremely low ammonia nitrogen concentrations that were within the safe range for aquaculture.

Figure 2b shows the variation in nitrite nitrogen concentration in aquaculture barrels over time (0–12 hours) at different circulation rates (1.5, 3, or 4.5 m^3^/h) under the initial environment of fresh water in the circulating water aquaculture system with a feeding amount of 100 g. When the circulation rate was adjusted to 1.5 and 3.0 m^3^/h, the water treatment system of the water circulation pump started to work effectively. The nitroso nitrogen concentration in breeding barrels 1 and 2 increased synchronously over time and then decreased. The initial values of nitroso nitrogen concentration in breeding barrels 1 and 2 at the 0^th^ hour were 0.04 and 0.05mg/Land 0.05 and 0.05mg/L), respectively, indicating that the performance of the biofilter remained relatively stable. The initial conditions set in the experiment were within the error range. There was no significant difference between the values recorded. When the circulation rate was 1.5 or 3.0 m^3^/h, the maximum values (0.09, 0.10mg/L and 0.07, 0.07 mg/L) were reached in breeding barrels 1 and 2 at the 8^th^ and 6^th^ hours, respectively. This indicates that the feces produced by the feeding offspring of *S. wangchiachii* entered the water body to accumulate and enhance nitroso nitrogen concentrations. As the circulation rate increased, the removal rate of nitroso nitrogen by the biofilter also increased, greatly shortening the time for nitroso nitrogen accumulation. The water quality in the breeding bucket was maintained in good condition at all times. When the circulation rate was 4.5 m^3^/h, the water treatment system of the water circulation pump initiated its function. The concentration of nitroso nitrogen in buckets 1 and 2 decreased synchronously over time, then increased, and then decreased again. After four hours, buckets 1 and 2 reached the minimum values of 0.02 and 0.03 mg/L, respectively. This indicated that the increase in circulation rate directly affects the removal efficiency of the biofilter. By applying water circulation at a suitable rate, the concentration of harmful substances in the bucket can be quickly reduced, ensuring the healthy growth of cultured fish. At different circulation rates (1.5, 3, or 4.5 m^3^/h) at the 12^th^ hour, the concentration of nitroso nitrogen in breeding barrels 1 and 2 gradually decreased, indicating that the biofilter had achieved a certain removal.

### 3.2 The effect of different circulation rates on the water quality of the aquaculture bucket at a feed rate of 300 g

Figure 3a shows the variation in ammonia nitrogen concentration in aquaculture barrels over time (0–12 hours) at different circulation rates (1.5, 3, or 4.5 m^3^/h) under the initial environmental conditions of 300 g of feed and fresh water in the circulating water aquaculture system. When maintaining the circulation rate of 1.5, 3.0, or 4.5 m^3^/h, the water treatment system of the circulating water pump worked effectively. The ammonia nitrogen concentration of breeding barrels 1 and 2 increased synchronously and then decreased over time. The initial values of ammonia nitrogen concentration of breeding barrels 1 and 2 at the 0^th^ hour were 0.31 and 0.37 mg/L; 0.34 and 0.36 mg/L; and 0.33 and 0.34 mg/L), respectively, indicating that the water treatment efficiency of the biofilter was relatively stable, and the initial conditions set in the experiment were within the error range. There was no significant difference between them. At the 4^th^ hour, the maximum values of 0.73 and 0.73 mg/L; 0.58 and 0.55 mg/L; and 0.48 and 0.49 mg/L were reached in breeding buckets 1 and 2, respectively. This indicated that the feces and residual bait produced by the feeding offspring of *S. wangchiachii* quickly accumulated ammonia nitrogen in the water body, and the biological filter only had a certain filtering effect. After four hours, the decrease in ammonia nitrogen concentration in breeding buckets 1 and 2 indicated that the effectiveness of biofilter filtration in removing harmful substances became gradually evident. When keeping the circulation rate of 1.5, 3.0, or 4.5 m^3^/h, the ammonia nitrogen concentrations in breeding barrels 1 and 2 reached extremely low values (0.47 and 0.43 mg/L; 0.33 and 0.31 mg/L; and 0.33, 0.30 mg/L) at the 12^th^ hour, respectively. At this time, the effectiveness of the biofilter reached the optimal removal effect, maintaining extremely low ammonia nitrogen concentration that is within the safe range of breeding.

**Fig. 3.**
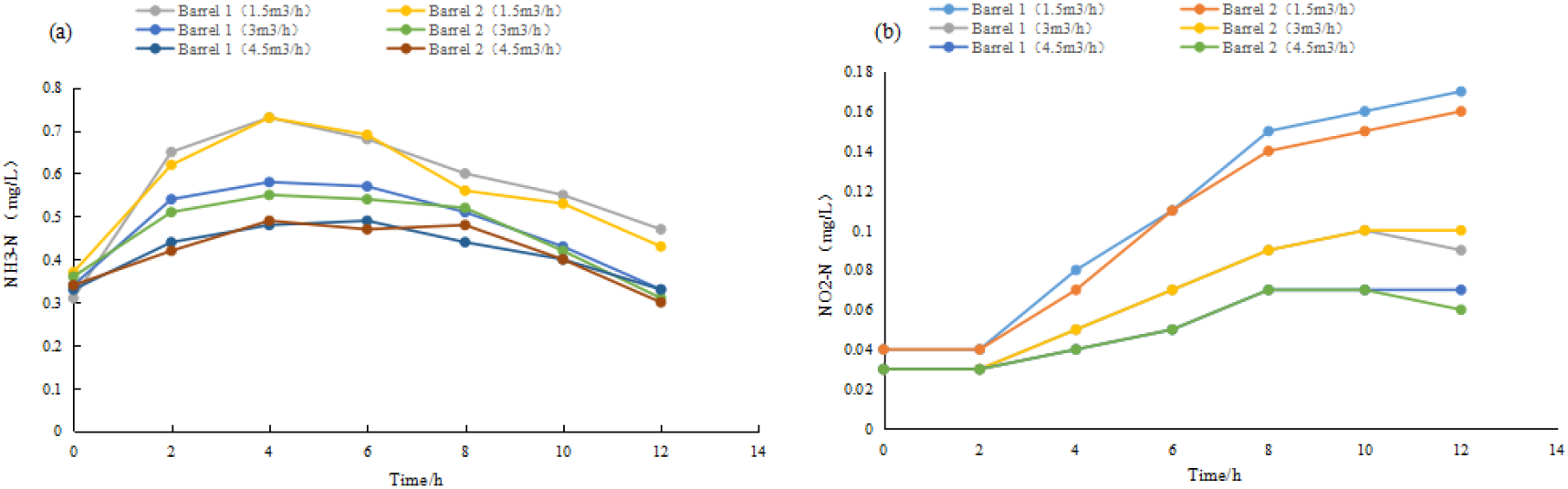
Changes in ammonia nitrogen and nitroso nitrogen in breeding barrels with different cycle rates over time with 300 g of feed.

Figure 3b shows the variation in nitrite nitrogen concentration in aquaculture barrels over time (0–12 hours) at different circulation rates (1.5, 3.0, or 4.5 m^3^/h) under the initial environment of 300 g of feed and fresh water in the circulating water aquaculture system. When keeping the circulation rate of 3.0 or 4.5 m^3^/h, the water treatment system of the circulating water pump worked effectively, and the concentration of nitroso nitrogen in aquaculture buckets 1 and 2 showed a synchronous increase and then a decreasing trend over time. The initial values of nitroso nitrogen concentrations in breeding barrels 1 and 2 at the 0^th^ hour were 0.03 and 0.03 mg/L and 0.03 and 0.03 mg/L), respectively, indicating that the performance of the biofilter was relatively stable. The initial conditions set in the experiment were within the error range, and there was no significant difference between the measurements. At the 10^th^ hour, buckets 1 and 2 reached maximum values (0.10 and 0.10 mg/L and 0.07 and 0.07 mg/L), respectively, indicating that the feces produced by the feeding offspring of *S. wangchiachii* accumulated nitroso nitrogen concentration in the water body, leading to an increase in nitroso nitrogen concentrations. The removal rate of nitroso nitrogen by the biofilter also increased with an increase in the circulation rate. The relatively large feeding amount resulted in no significant effect of the biofilter on removal. At the 12^th^ hour, the concentration of nitroso nitrogen in the breeding buckets 1 and 2 began to slowly decrease, but with no significant effect. When the circulation rate was 1.5 m^3^/h, the water treatment system of the circulating water pump started to work effectively. The nitroso nitrogen concentration of the breeding barrels 1 and 2 showed a synchronous upward trend over time. The nitroso nitrogen concentration of the breeding barrels 1 and 2 did not show an upward trend and maintained a certain value from 0 to 2 hours, indicating that the accumulation rate of nitroso nitrogen concentration was slow in the early stages of feeding. When the circulation rate was extremely low, the biofilter seemed to play a significant role in improving the water quality in the breeding barrels. The concentration of nitroso nitrogen in buckets 1 and bucket 2 gradually increased from 2 to 8 hours, indicating an increase in the excretion rate of metabolic ammonia and rapid accumulation of ammonia nitrogen in the dissolved water of the *S. wangchiachii*. The slow increase in nitroso nitrogen concentration in breeding barrels 1 and 2 from 8 to 12 hours indicated a decrease in the metabolic ammonia excretion rate of *S. wangchiachii* and the accumulation of water treatment efficiency of the biofilters. Although the concentration of nitroso nitrogen could not be reduced in a timely manner, it provided a scientific reference for further optimization of experimental conditions.

### 3.3 The effect of different circulation rates on the water quality of the aquaculture bucket at a feed rate of 500 g

Figure 4a shows the changes in ammonia nitrogen concentration in aquaculture barrels over time (0–12 hours) at different circulation rates (1.5, 3, or 4.5 m^3^/h) under the initial environmental conditions of 500 g of feed and fresh water in the circulating water aquaculture system. When adjusting and keeping the circulation rate of 1.5, 3.0, or 4.5 m^3^/h, the water treatment system of the circulating water pump began to work effectively. The ammonia nitrogen concentration in breeding barrels 1 and 2 showed a synchronous upward trend over time. The initial values of ammonia nitrogen concentration in breeding barrels 1 and 2 at the 0^th^ hour were 0.33 and 0.36 mg/L; 0.35 and 0.34 mg/L; and 0.36 and 0.39 mg/L, respectively, indicating that the water treatment efficiency of the biofilter was relatively stable, and the initial conditions set in the experiment were within the error range. There was no significant difference between the measurements. The rapid increase in ammonia nitrogen concentration in aquaculture buckets 1 and 2 from 0 to 4 hours indicated that the feces and residual bait produced by the feeding offspring of *S. wangchiachii* quickly accumulated ammonia nitrogen in the water, and the filtration effect was not significant. The ammonia nitrogen concentration of aquaculture buckets 1 and 2 still increased but slowed down after 4–12 hours, indicating that the effectiveness of biofilter filtration in removing harmful substances exceeded the load, and the aquaculture water could gradually deteriorate, affecting the healthy growth of fish. At the 12^th^ hour, the ammonia nitrogen concentrations in buckets 1 and 2 reached extremely low values (1.12 and 1.14 mg/L; 1.05 and 1.01 mg/L; and 0.93 and 0.90 mg/L), respectively. At this time, the efficiency of the biofilter was overloaded, and it could not maintain extremely low ammonia nitrogen concentrations and be within the safe range of aquaculture. The accumulation of ammonia nitrogen in the later stage was highly likely to experimentally affect aquaculture.

**Fig. 4.**
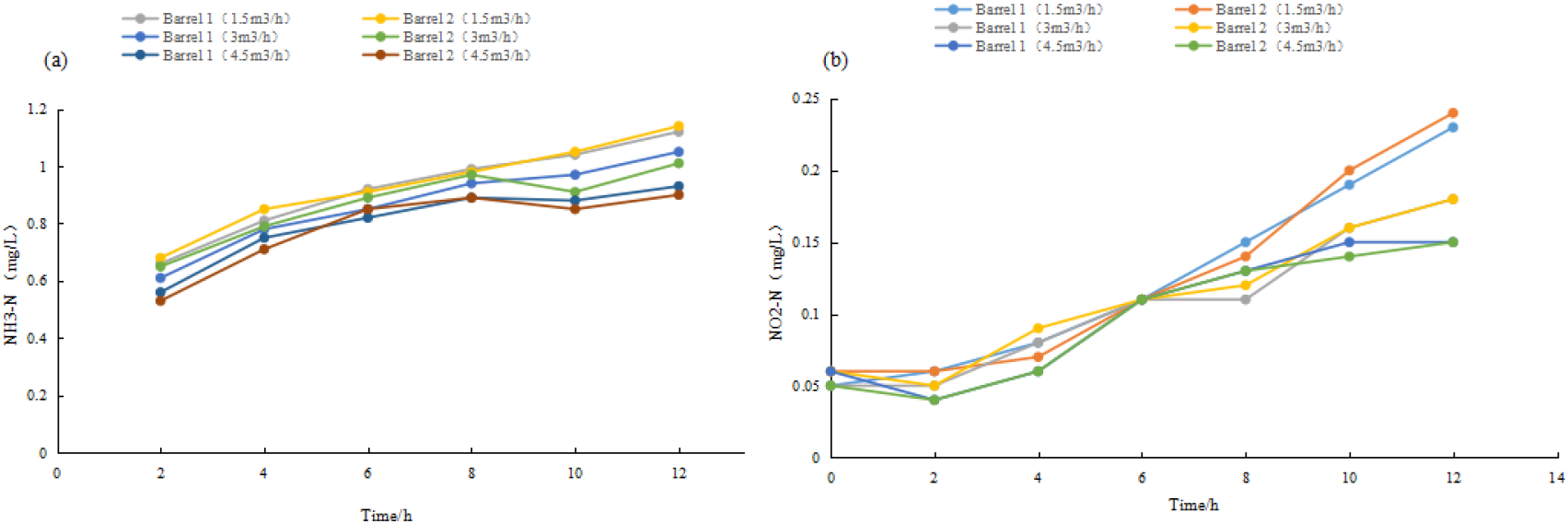
Changes in ammonia nitrogen and nitroso nitrogen in breeding barrels with different cycle rates over time with 500 g of feed.

Figure 4b shows the variation in nitrite nitrogen concentration in aquaculture barrels over time (0–12 hours) at different circulation rates (1.5, 3.0, or 4.5 m^3^/h) under the initial environment of 300 g of feed and fresh water in the circulating water aquaculture system. When maintaining a circulation rate of 3.0 or 4.5 m^3^/h, the water treatment system of the circulation water pump worked effectively, and the nitroso nitrogen concentration of breeding barrels 1 and 2 showed a synchronous and slow decrease in a certain value over time, followed by a rapid increase. The initial values in the nitroso nitrogen concentrations in breeding barrels 1 and 2 at the 0^th^ hour were 0.05 and 0.06 mg/L and 0.06 and 0.05 mg/L, respectively, indicating that the performance of the biofilter was relatively stable. The initial conditions set in the experiment were within the error range, and there was no significant difference between them. The slow decrease in nitroso nitrogen concentration in the aquaculture buckets 1 and 2 from 0 to 2 hours indicated a slow accumulation rate of nitroso nitrogen concentration in the early stages of feeding. When the circulation rate was extremely low, biofilters could also play a significant role in improving the water environment of the aquaculture buckets. The rapid increase in nitroso nitrogen concentration in buckets 1 and 2 after 2–12 hours of cultivation indicates that the feces produced by the feeding offspring of *S. wangchiachii* enter the water body to accumulate nitroso nitrogen concentration, leading to an increase in nitroso nitrogen concentration. The rate of removal of nitroso nitrogen by the biofilter also increases with an increase in the circulation rate, but the relatively large feeding amount causes the biofilter to have no significant effect on removal. When the circulation rate was 1.5 m^3^/h, the water treatment system of the circulating water pump worked and the water treatment system began to take effect. The concentration of nitroso nitrogen in buckets 1 and 2 increased rapidly over time from 0 to 12 hours. This indicated that the metabolic ammonia excretion rate of *S. wangchiachii* increased, and the residual feed dissolved in water and quickly led to the accumulation of ammonia nitrogen. The biological filter basically did not reflect the water treatment efficiency. Further improvement is needed in the experimental conditions to achieve appropriate breeding conditions.

### 3.4 The effect of biofilters on removing ammonia nitrogen and nitroso nitrogen under different circulation rates at a feeding rate of 100 g

As shown in Figure 5a, with the feeding amount of 100 g and the initial environmental conditions of fresh water in the circulating aquaculture system, the trend curves of the ammonia nitrogen removal rate with the biofilter over time (0– 12h) at different circulation rates (1.5, 3, or 4.5 m^3^/h) presented a slow increase, followed by a gradual decrease. When maintaining the circulation rates of 1.5, 3.0, or 4.5 m^3^/h, the ranges of change in the removal rate of ammonia nitrogen by the biofilter were 29.5–54.2%, 25.6–43.8%, and 20.0–34.5%, respectively, indicating that the removal performance of the biofilter was relatively stable. When the circulation rate increased, the removal rate of ammonia nitrogen by the biofilter decreased. According to the variation in the polynomial trend curves under different cycle rates, the removal rate of ammonia nitrogen by the biofilter reached a corresponding maximum value within 4–6 hours, indicating that the feces produced by the feeding offspring of *S. wangchiachii* entered the water body to enhance ammonia nitrogen concentration; the ammonia nitrogen removal rate increased accordingly. Under different circulation rates (1.5, 3, or 4.5 m^3^/h), the average removal rates of ammonia nitrogen by biofilters from 0 to 12 hours were 40.4, 36.8, and 27.7%, respectively, indicating that hydraulic retention time was a major factor affecting the water treatment performance of the biofilters.

**Fig. 5.**
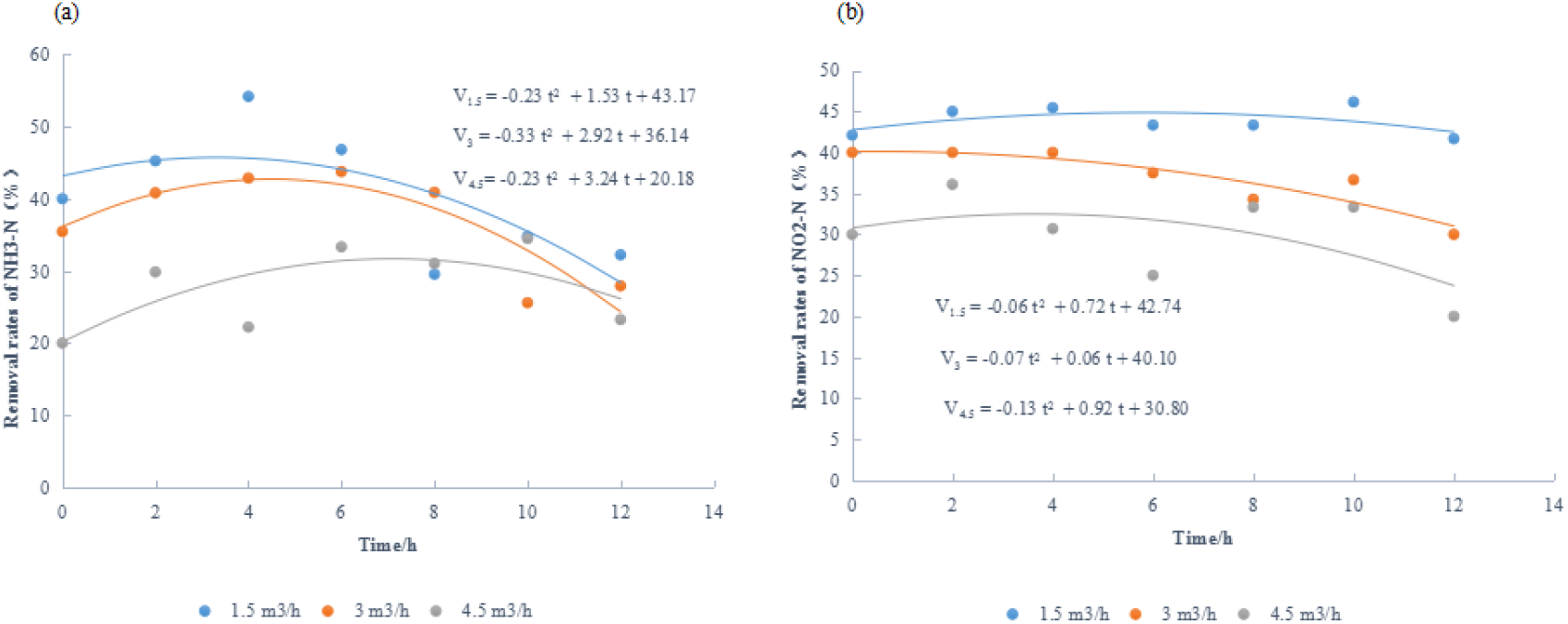
Removal rates of ammonia nitrogen and nitroso nitrogen by biofilter under different cycle rates with 100 g of feed.

As shown in Figure 5b, with the feeding amount of 100 g and the initial environment of fresh water in the circulating aquaculture system, the trend curves of the removal rate of nitroso nitrogen by biofilters at different circulation rates (1.5, 3, or 4.5 m^3^/h) over time (0–12 hours) showed a slow decrease. When maintaining the circulation rate of 1.5, 3.0, or 4.5 m^3^/h, the ranges of changes in the removal rate of nitrite nitrogen by the biofilter were 41.7–46.2%, 30.0–40.0%, and 20.0–36.1%, respectively, indicating that the removal performance of the biofilter was relatively stable. When the circulation rate increased, the removal rate of nitrite nitrogen by the biofilter decreased: According to the variation in polynomial trend curves with different cycle rates, the fluctuation in the removal efficiency of nitroso nitrogen by biofilters was small within 0–12 hours, indicating that the growth of sub-digestive bacteria in biofilters was relatively mature and their performance was stable. Under the different circulation rates (1.5, 3, or 4.5 m^3^/h), the average removal rates of nitroso nitrogen by biofilters from 0 to 12 hours were 43.9, 36.9, and 29.8%, respectively, indicating an inverse relationship between hydraulic retention time and biofilter removal efficiency.

### 3.5 The effect of biofilters on removing ammonia nitrogen and nitroso nitrogen under different circulation rates at a feeding rate of 300 g

Figure 6a shows the initial environmental conditions of fresh water and those under the feeding amount of 300 g in the circulating aquaculture system. It also shows trend curves of the biofilter’s ammonia nitrogen removal rate (0–12h) at different circulation rates (1.5, 3, or 4.5 m^3^/h); there was a slow increase followed by a gradual decrease. When regulating the circulation rate at 1.5, 3.0, or 4.5 m^3^/h, the range of changes in the removal rate of ammonia nitrogen by the biofilter were 34.6–60.3%, 27.7–47.1%, and 20.0–45.2%, respectively, indicating that the removal performance of the biofilter was relatively stable. When the circulation rate increased, the removal rate of ammonia nitrogen by the biofilter decreased. According to the variation in polynomial trend curves under different cycle rates, the removal rate of ammonia nitrogen by the biofilter reached the corresponding maximum value within 4–8 hours, indicating that the feces produced by the feeding offspring of *S. wangchiachii* enter the water body to accumulate ammonia nitrogen, leading to an increase in its concentration; the ammonia nitrogen removal rate increases accordingly. Under different circulation rates (1.5, 3, or 4.5 m^3^/h), the average removal rates of ammonia nitrogen by biofilters from 0 to 12 hours were 42.3, 37.9, and 33.6%, respectively, indicating that hydraulic retention time was a major factor affecting the water treatment performance of biofilters.

**Fig. 6.**
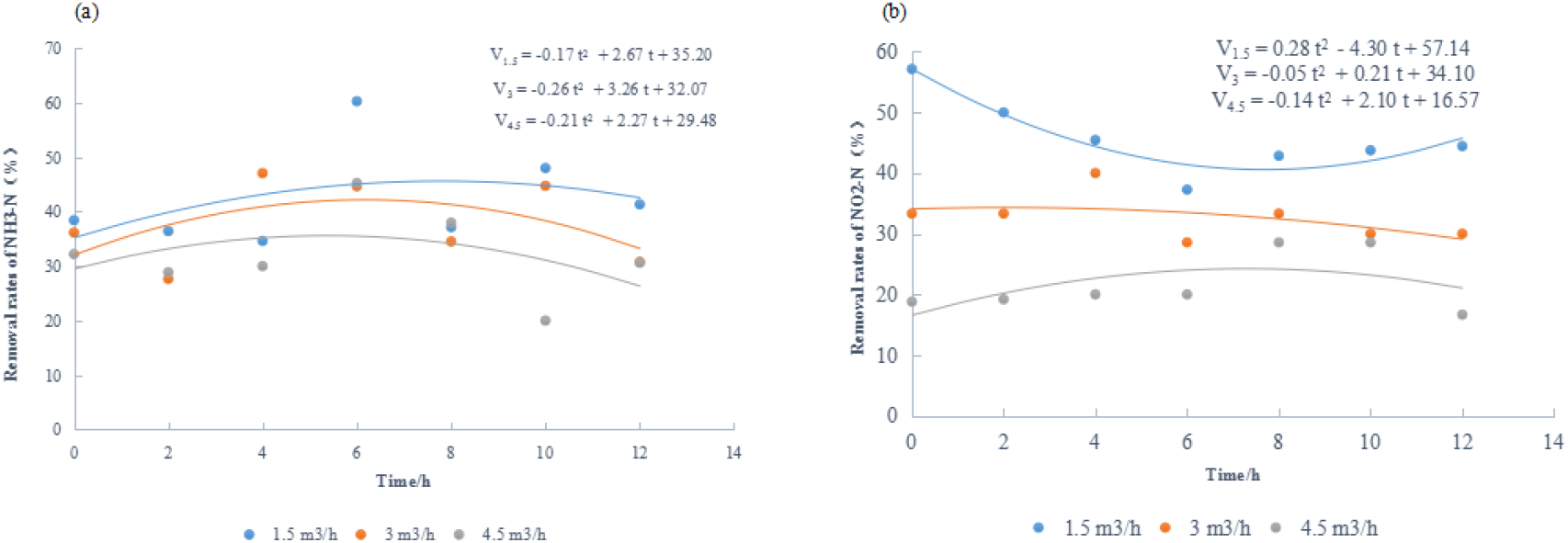
Removal rates of ammonia nitrogen and nitroso nitrogen by biofilter under different cycle rates with 300 g of feed.

As shown in Figure 6b, under the feeding amount of 300 g and the initial environment of fresh water in the circulating aquaculture system, the trend curves of the removal rate of nitroso nitrogen by biofilters at different circulation rates (3 or 4.5 m^3^/h) (0–12h) showed a slow increase, followed by a decrease. The trend curves of the removal rate of nitroso nitrogen by biofilters with a circulation rate of 1.5 m^3^/h (0– 12h) showed a slow decrease, followed by an increase. When regulating the circulation rate at 1.5, 3.0, or 4.5 m^3^/h, the ranges of change in the removal rate of nitrite nitrogen by the biofilter were 37.3–57.1%, (28.6–40.0%, and 16.7–28.6%, respectively, indicating that the removal performance of the biofilter was relatively stable. When the circulation rate increased, the ranges of the removal rate of nitrite nitrogen by the biofilter decreased. According to the variation in polynomial trend curves under different cycle rates, the fluctuation in the removal efficiency of nitroso nitrogen by biofilters was small within 0–12 hours, indicating that the growth of sub-digestive bacteria in biofilters was relatively strong, and their performance was stable. Under different circulation rates (1.5, 3, or 4.5 m^3^/h), the average removal rates of nitroso nitrogen by biofilters from 0 to 12 hours were 45.8, 32.7, and 21.7, respectively, indicating an inverse relationship between the hydraulic retention time and biofilter removal efficiency.

### 3.6 The effect of biofilters on removing ammonia nitrogen and nitroso nitrogen under different circulation rates at a feeding rate of 500 g

Figure 7a shows that the trends of the ammonia nitrogen removal rate of biofilters at different circulation rates (1.5, 3, or 4.5 m^3^/h) over time (0–12 hours) presented a slow increase, followed by a gradual decrease, under the initial environmental conditions of 500 g of feed and fresh water. When regulating the circulation rate at 1.5, 3.0, or 4.5 m^3^/h, the ranges of change in the removal rate of ammonia nitrogen by the biofilter were 42.5–63.5%, 32.3–50.0%, and 28.3–48.0%, respectively, indicating that the performance of the biofilter was relatively stable. When the circulation rate increased, the removal rate of ammonia nitrogen by the biofilter decreased. According to the variation in polynomial trend curves under different cycle rates, the fluctuation range of ammonia nitrogen removal in the biofilter was small, i.e., from 0 to 12 hours, indicating that the biofilter hosted a stable growth of ammonifying bacteria and had a strong ability to absorb harmful substances. Under different circulation rates (1.5, 3, or 4.5 m^3^/h), the average removal rates of ammonia nitrogen by biofilters from 0 to 12 hours were 53.5, 42.6, and 35.7%, respectively, indicating that hydraulic retention time was a major factor affecting the water treatment performance of biofilters.

**Fig. 7.**
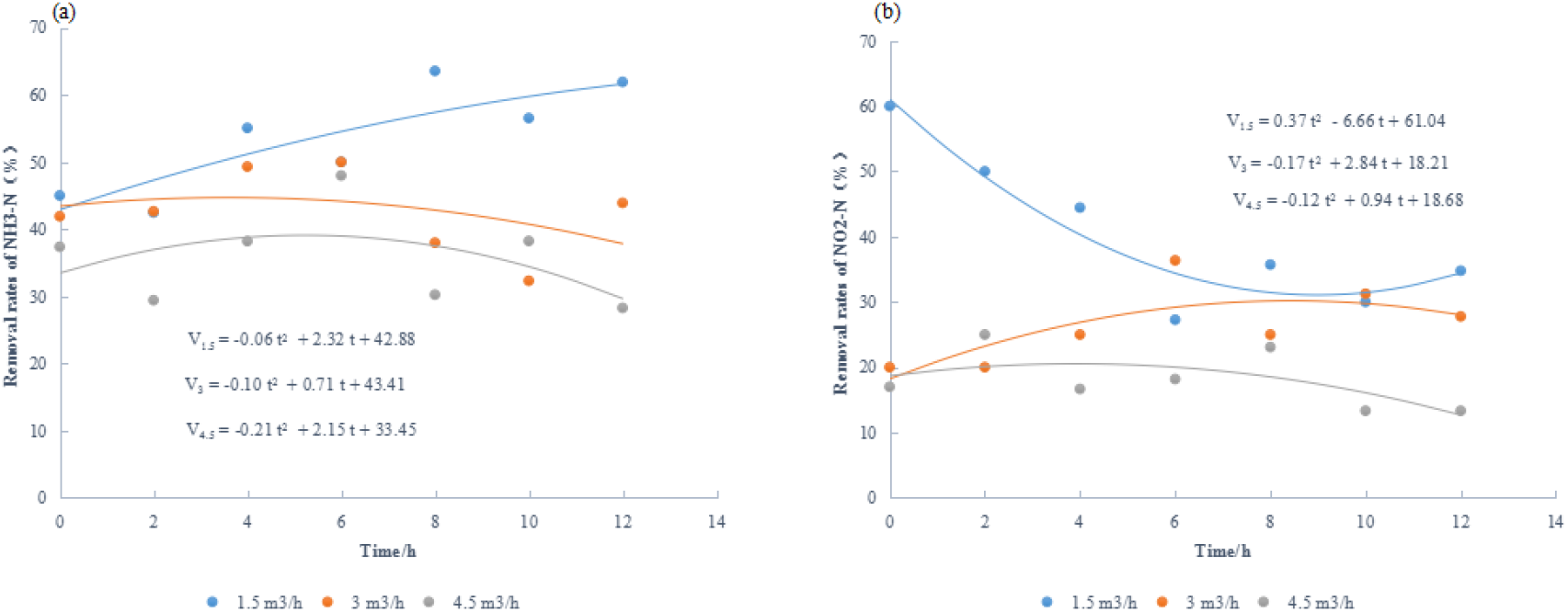
Removal rates of ammonia nitrogen and nitroso nitrogen by biofilter under different cycle rates with 500 g of feed.

Figure 7b shows the trends of the removal rate of nitroso nitrogen by biofilters at different circulation rates (3 or 4.5 m^3^/h) over time (0–12h). A slow increase is followed by a decrease under the initial environmental conditions of 500 g of feed and fresh water in the recirculating aquaculture system. The trend of the removal rate of nitroso nitrogen by biofilters with a circulation rate of 1.5 m^3^/h over time (0–12h) shows a slow decrease followed by an increase. When regulating the circulation rate at 1.5, 3.0, or 4.5 m^3^/h, the ranges of change in the removal rate of nitroso nitrogen by the biofilter were 27.3–60.0%, 20.0–36.4%, and 13.3–25.0%, respectively, indicating that the removal performance of the biofilter was relatively stable. When the circulation rate increased, the value of the removal rate of nitroso nitrogen by the biofilter decreased. According to the variation in polynomial trend curves under different cycle rates, the fluctuation in the removal efficiency of nitroso nitrogen by biofilters was small within 0–12 hours, indicating that the growth of sub-digestive bacteria in the biofilters was relatively mature and their performance was stable. Under different circulation rates (1.5, 3, or 4.5 m^3^/h), the average removal rates of nitroso nitrogen by biofilters from 0 to 12 hours were 40.3, 26.5, and 18.1%, respectively, indicating an inverse relationship between hydraulic retention time and biofilter removal efficiency.

## 4. Discussion

### 4.1 Changes in water quality of aquaculture barrels during static water aquaculture with different feeding rates

In some studies on the regulation of ammonia production or release in aquaculture experiments, Wang (Wang *et al*., 2009) showed that the relative concentration of ammonia nitrogen emitted from the semi-smooth tongue sole reached its peak value 3 to 6 hours after feeding. Wang (Wang *et al*., 2001) found that within a certain range of feed intake, the ammonia excretion rate of *Ruditapes Philippinarum* increases with increased feed intake, leading to higher ammonia nitrogen concentration in aquaculture water bodies. Other studies found that the ammonia excretion rate of meretrix increases with the increase in feed concentration (e.g., for *Phaeodactylum tricornutum* and *Isochrysis galbana*), resulting in a rapid increase in ammonia nitrogen concentration in aquaculture water (Tang *et al*., 2005). The ammonia excretion rate for sea cucumber (*Apostichopus japonicus*) increases with the increase in protein content in the feed. The above analysis indicates that the concentration of ammonia nitrogen, which is a water quality indicator, will gradually increase after aquaculture organisms feed (Zonghe *et al*., 2020). This experiment tests three feeding rates in the same water environment. In the still-water aquaculture mode, as the feeding rate increases, the ammonia nitrogen concentration in the aquaculture bucket gradually increases, which is consistent with the above research results. At the same time, there is no significant correlation between the ammonia nitrogen concentrations in aquaculture buckets 1 and 2, indicating that the overall control of the quality of the circulating water aquaculture system was excellent, and the system’s aquatic environment was stable and consistent, promoting the rapid growth of fish. The feeding amount was positively correlated with the ammonia nitrogen concentration in the aquaculture bucket (*P*<0.05), indicating that feed is its main source after feeding. The specific analysis of water quality changes in aquaculture water bodies indicated that an increase in the intake of aquaculture fish promotes their metabolic activity, maintains their healthy growth and normal physiological activities, and decomposes excessive feed. Therefore, the discharge of metabolic waste, such as ammonia, and the concentration of harmful substances into aquaculture water bodies, is also increasing.

### 4.2 Changes in water quality of aquaculture barrels in a circulating water system with different circulation rates under a certain feeding amount

The main characteristics of the new model of industrialized aquaculture with circulating water are water conservation, efficient and precise aquaculture, and intelligent and scientifically-informed aquaculture management (Nielsen, 2011; Shitu *et al*., 2022). The water circulation rate of the recirculating aquaculture system not only controls the energy demand of the aquaculture system, but also involves the physiological and growth status of the aquaculture objects. Accurately finding the appropriate water circulation rate and mastering the water flow speed in a given aquaculture system can help achieve the lowest energy consumption and excellent biological water treatment (Song *et al*., 2013), which is an important technical issue in industrial recirculating aquaculture models. According to the factory-designed circulating water aquaculture system discussed in the previous section, a set of systems has been designed and built. In the later stage, system debugging and practical experiments are needed to demonstrate the stability and positive impact on the aquaculture water environment through water quality control. The issue of energy consumption is key to constraining the industrialized circulating water aquaculture model; in addition, the cost of aquaculture electricity accounts for the majority of the entire expenditure. A precise management system to reduce energy consumption is a good way to improve the profits of industrialized circulating water aquaculture. The design of this experiment aimed to understand the relationship between feeding amount and circulation rate. Precise control of the circulation rate can enhance the biofilter treatment of aquaculture water. At the same time, reducing the electricity consumption of circulating water pumps can reduce the cost of aquaculture, providing a certain economic and scientific basis for the promotion of industrialized circulating water aquaculture models.

After the concentration of harmful substances, such as ammonia nitrogen, in the aquatic environment of the aquaculture reaches a certain value (Wantasen *et al*., 2020), non-ionized ammonia is prone to entering the bodies of the aquatic environment through the cell membrane, leading to physiological imbalance and disorder, leading to tissue hypoxia and toxicity (Li *et al*., 2007). At the same time, a certain concentration of nitrite can inhibit and affect the growth, development, and reproduction of aquaculture organisms, which can damage or even kill the oxygen-carrying capacity of hemocyanins (Chand & Sahoo, 2006). In the industrial recirculating aquaculture system, ammonia nitrogen in the aquaculture water is generally oxidized to nitrate by nitrifying bacteria in the biological filter (Zou *et al*., 2018). In addition, the utilization rate of feed protein by the aquaculture fish is generally only 20 to 25%, while the remaining feed is discharged into the aquaculture water in the form of residual bait and feces. This can lead to eutrophication in the aquaculture water and readily consume dissolved oxygen that aquaculture organisms rely on for survival (Crab *et al*., 2007). Therefore, improving the water circulation rate and enhancing the efficacy of water treatment by biological filters is at the core of a successful industrialized circulating water aquaculture. In this experimental study, three gradient feeding rates (100, 300, and 500 g) and the same initial aquatic environment were set to simulate different feeding rates at different stages of fish growth. The circulation rates were then regulated to accurately manage the aquaculture system. Different conditions (1.5, 3, or 4.5 m^3^/h) were set for the circulation rate, and as the circulation rate increased, the water treatment effect of the biofilter became significant. As the circulation rate increased, the concentration of ammonia nitrogen and nitroso nitrogen in the breeding bucket was rapidly removed until it reached a lower level with the same amount of feed additives. Conclusively, when the feeding amount was 100 and 300 g, the circulation rates were set to 1.5 and 3 m^3^/h, respectively. Currently, the circulating water aquaculture method is the best management method for aquaculture that achieves precise aquaculture and cost savings. When the feeding amount is 500 g, and the circulation rates are set at 1.5, 3, or 4.5 m^3^/h, although the biofilter may have a certain removal effect, it is not able to suppress the increase in ammonia nitrogen and nitroso nitrogen. A higher circulation rate is needed to regulate aquaculture water quality, which may also exceed the filtration load of existing biological filters. Therefore, the overload design of biological filters and the load of recirculating aquaculture are key issues in the design of recirculating aquaculture systems.

### 4.3 Water treatment efficiency of biological filters in recirculating aquaculture systems

In this experimental study, the percentages of ammonia nitrogen and nitroso nitrogen removed by the biofilter were calculated to test the water treatment efficiency of the biofilter. Biofilters are developed by pre-culturing biofilms and conducting experiments on aquaculture water treatment. The hydraulic retention time is a key factor that directly affects the removal rate of biofilters (Rodziewicz *et al*., 2019). When the hydraulic retention time is small, the removal efficiency of the biofilter is large, but the water circulation rate of the aquaculture system is also relatively small, which directly affects the system’s water quality. When the hydraulic retention time is long, the removal efficiency of the biofilter is relatively low, but the water circulation rate of the system is very high. Finally, in this setting, the concentration of harmful substances in the stable aquaculture water body falls within a safe range.

## 5. Conclusions

It is a good approach to experimentally find a suitable circulation rate for industrial circulating water aquaculture models using different feeding rates. This can help achieve precise management and reduce system energy consumption to improve the benefits of circulating water aquaculture. Feed is the main source of nitrogen in aquaculture water, which comes mainly through fish excretion and residual bait added to the water. As the amount of feed increases, the ammonia nitrogen concentration in the aquaculture water gradually increases, and it becomes necessary to reduce it through treatment with biological filters. Therefore, biofilters are among the core components of recirculating aquaculture. Hydraulic retention time is a key factor that directly affects the removal efficiency of biofilters; it is positively correlated with the removal rate.

## Declarations

### Ethical Approval Statement

The animal subjects used in the study were fishes, which are vertebrates and exempt from this requirement. All experiments were performed according to the Experimental Animal Management Law of China and approved by the Animal Ethics Committee of Xichang University. The study is reported in accordance with ARRIVE guidelines.

### Informed consent

Not applicable

### Funding

This work was supported by PhD Start-up Project of Xichang University (YBZ202240) and Wuzhou City Scientific Research and Technology Development Plan Project (2022E02054). The funders had role in the study design, data collection and analysis, decision to publish, or preparation of the manuscript.

### Contributions

QL Cai and JM Zhou conceived and designed the study. QL Cai and JM Zhou performed tissue sample preparations from experimental animals. All other experiments were performed and analyzed by JM Zhou; JM Zhou, QL Cai and KL Zhou were also involved in the experimental work. JM Zhou wrote the manuscript with support from all authors. All authors read and approved the final manuscript.

### Competing interests

The authors declare that they have no competing interests.

### Editorial certificate

The manuscript below was edited for correct English language usage, grammar, punctuation and spelling by qualified native English speaking editors at Charlesworth Author Services.

### Data availability

During the experiment, the experimental errors have been reduced as much as possible and the variables have been well controlled. The experimental data in this paper will not be published in other papers. The author declares to be responsible for the reliability of the data.

